# Sex peptide receptor is not required for refractoriness to remating or induction of egg laying in *Aedes aegypti*

**DOI:** 10.1101/2023.06.28.546954

**Authors:** I. Alexandra Amaro, Margot P. Wohl, Sylvie Pitcher, Catalina Alfonso-Parra, Frank W. Avila, Andrew S. Paige, Michelle Helinski, Laura B. Duvall, Laura C. Harrington, Mariana F. Wolfner, Conor J. McMeniman

## Abstract

Across diverse insect taxa, the behavior and physiology of females dramatically changes after mating – processes largely triggered by the transfer of seminal proteins from their mates. In the vinegar fly *Drosophila melanogaster*, the seminal protein sex peptide (SP) decreases the likelihood of female flies remating and causes additional behavioral and physiological changes that promote fertility including increasing egg production. Although SP is only found in the *Drosophila* genus, its receptor, sex peptide receptor (SPR), is the widely-conserved myoinhibitory peptide (MIP) receptor. To test the functional role of SPR in mediating post-mating responses in a non-*Drosophila* dipteran, we generated two independent *Spr*-knockout alleles in the yellow fever mosquito *Aedes aegypti*. Although SPR is needed for post-mating responses in *Drosophila* and the cotton bollworm *Helicoverpa armigera*, *Spr* mutant *Ae. aegypti* show completely normal post-mating decreases in remating propensity and increases in egg laying. In addition, injection of synthetic SP or accessory gland homogenate from *D. melanogaster* into virgin female mosquitoes did not elicit these post-mating responses. Our results indicate that *Spr* is not required for these canonical post-mating responses in *Ae. aegypti*, indicating that unknown signaling pathways are likely responsible for these behavioral switches in this disease vector.

## Introduction

Female insects undergo major changes after mating that alter their behavior and physiology, ultimately contributing to successful reproduction. Much of our understanding of these changes comes from work in *Drosophila melanogaster*, where post-mating changes are primarily initiated by seminal fluid proteins derived from male reproductive organs and transferred to females within the ejaculate (reviewed in AVILA *ET AL.* 2011; WIGBY *ET AL.* 2020). Many post-mating changes in *D. melanogaster* are induced by a 36 amino acid seminal “sex peptide” (SP); its most emblematic effects are decreasing the female’s receptivity to remating and increasing her egg production (CHEN *et al*. 1988; CHAPMAN *et al*. 2003; LIU AND KUBLI 2003). SP exerts many of its effects through a G protein-coupled receptor, SPR (the sex peptide receptor) that is expressed in neurons as well as in reproductive tract tissues (YAPICI *et al*. 2008; HASEMEYER *et al*. 2009; YANG *et al*. 2009).

Impairment or loss of SPR in *D. melanogaster* leads to a reduction in post-mating egg laying, and to refractoriness to remating (YAPICI *et al*. 2008), as well as impairing additional post-mating changes in sleep patterns (GARBE *et al*. 2016), sperm release (AVILA *et al*. 2015), long-term memory (SCHEUNEMANN *et al*. 2019), and gut growth (WHITE *et al*. 2021). These responses have been shown to be integrated largely through the Sex-Peptide sensory and abdominal ganglion neurons (SPSN-SAG network) (FENG *et al*. 2014; WANG *et al*. 2020; WANG *et al*. 2021; reviewed in OKAMOTO AND WATANABE 2022). Consistent with the rapid evolution seen for many reproductive proteins (e.g. CIVETTA AND SINGH 1998; SWANSON AND VACQUIER 2002; HAERTY *et al*. 2007), SP is only found in some species of *Drosophila* (TSUDA AND AIGAKI 2016). However, SPR is conserved across several *Drosophila* species (KIM *et al*. 2010) and other insects (YAPICI *et al*. 2008). This widespread sequence conservation likely reflects SPR’s role as a myoinhibitory peptide (MIP) receptor, which has been considered to be its ancestral function. It is suggested that SPR was hijacked or co-opted to a reproductive function in lineages where it acquired expression in the female reproductive tract (KIM *et al*. 2010; POELS *et al*. 2010; TSUDA AND AIGAKI 2016). Experiments aimed at investigating a reproductive role for MIPs suggest that they do not induce post-mating responses, as mating receptivity was not altered by either pan-neuronal knockdown of MIPs or injection of MIPs in *D. melanogaster* females (KIM *et al*. 2010).

SPR has also been found to be important for post-mating responses in some insects in addition to *Drosophila*. RNAi knockdown of SPR expression in olive fruit flies (*Bactrocera oleae*) and Oriental fruit flies (*Bactrocera dorsalis*) led to lower oviposition rates (ZHENG *et al*. 2015; GREGORIOU AND MATHIOPOULOS 2020). Additionally, mutation of the SPR gene in *B. dorsalis* resulted in a reduction in egg-laying ability, viability of eggs laid, and underdeveloped ovaries (CHEN *et al*. 2023). Injection of *D. melanogaster* SP into the cotton bollworm, *Helicoverpa armigera* (which has no obvious SP gene in its genome) suppressed sex pheromone production leading to decreased calling behaviors; these effects did not occur if SPR was knocked down simultaneous with SP injection, indicating SP function is through SPR (HANIN *et al*. 2012). Moreover, *H. armigera* females with little to no SPR laid significantly fewer eggs after mating and displayed altered pheromone calling behavior and remating rates after 24 and 48 h (HANIN *et al*. 2012; LIU *et al*. 2021). A reproductive role for SPR was also reported in the tobacco cutworm *(Spodoptera litura*): females with SPR knockdown laid fewer eggs than controls after injection of male accessory gland lysate and remained receptive to mating (LI *et al*. 2014).

Collectively, these studies indicate that SPR plays important post-mating roles in several insect taxa. However, the role of SPR in medically important mosquitoes, such as *Aedes aegypti*, has not been determined. If SPR is involved in inducing post-mating responses it could potentially be a target for mosquito reproductive control efforts (CATOR *et al*. 2021). RNASeq analyses of *Ae. aegypti* have identified SPR transcripts in neuronal and reproductive tissues (ALFONSO-PARRA *et al*. 2016; MATTHEWS *et al*. 2016), consistent with its expression patterns in those *Drosophila* species where it exerts a reproductive role (TSUDA *et al*. 2015). Although the *Ae. aegypti* genome does not contain a recognizable SP gene (our unpublished genome searches), neither does the genome of *H. armigera*, where SPR is necessary for inducing post-mating responses (HANIN *et al*. 2012; LIU *et al*. 2021).

To test the functional role of SPR in mediating *Ae. aegypti* post-mating responses, we employed CRISPR/Cas9 genome editing to generate two independent mutant alleles in the SPR gene. We tested female mutants for fertility and survival, as well as for the most salient reproductive phenotypes: post-mating egg laying and decreased mating receptivity, both of which are induced by seminal fluid proteins (CRAIG 1967; LEAHY AND CRAIG 1965; HELINSKI *et al*. 2012; VILLARREAL *et al*. 2018). Our data indicate that despite the sequence conservation of SPR, it does not play a detectable role in these post-mating changes in *Ae. aegypti*. Consistent with this conclusion, our injection of synthetic *Drosophila* SP, which can bind to and activate *Ae. aegypti* SPR *in vitro* (YAPICI *et al*. 2008; KIM *et al*. 2010) or of *Drosophila* male accessory gland extracts into virgin or gravid *Ae. aegypti* females gave no effect on these post-mating behaviors. The lack of effects of SPR knockout on post-mating egg-laying and receptivity in this dipteran, in contrast to its activity in the dipteran *D. melanogaster* and the lepidopteran species *H. armigera* and *S. litura*, suggests that SPR may have been repetitively co-opted for a reproductive function. Our results indicate that unknown signaling pathways are likely responsible for the post-mating switches governing long-term refractoriness to remating and induction of egg laying in *Ae. aegypti*.

## Materials and methods

Our groups generated *Spr* knockout alleles and characterized them independently. These mutant alleles, which were unique and in different genetic backgrounds, yielded the same results where compared. We refer to these alleles, generated at Cornell and Johns Hopkins respectively, as *Spr^Δ235^*and *Spr^ECFP^*.

### Mosquito rearing

*Spr^Δ235^*: The *Spr^Δ235^* NHEJ allele was generated in a Thai background (e.g. HELINSKI AND HARRINGTON 2011), and its controls and backcrosses were of that background. Our Thai colony of *Ae. aegypti* derived from field-collected mosquitoes (15°72’N, 101°75’E) maintained since 2009 with annual supplementation and a homozygous transgenic line with dsRed-labelled sperm (SMITH *et al*. 2007) were used. After vacuum-hatching eggs, larvae were reared under uniform conditions to ensure medium body size adults (HELINSKI AND HARRINGTON 2011). Mosquitoes were maintained at 28°C and 70% RH with a 10 h light:10 h dark cycle that included two hours of simulated dusk and dawn. Virgin males and females were obtained by separating pupae by sex prior to adult eclosion. All mosquitoes were maintained on 10% w/v sucrose. Biological replicates were derived from independently hatched cohorts.

*Spr^ECFP^*: The *Spr^ECFP^* HDR allele was generated in the *LVPib12* strain (NENE *et al*. 2007), which also served as the genetic background for assays with this mutant allele. Mosquitoes were maintained with a 12 h light:dark photoperiod at 27°C and 80% relative humidity using a standardized rearing protocol (WOHL AND MCMENIMAN 2023a). Adult mosquitoes were provided constant access to a 10% w/v sucrose solution and virgins were isolated as pupae and sexed within 12 h of emergence. The *Exu-Cas9* strain (marked with Opie2-dsRed) expressing *Cas9* under the maternal germline promoter *exuperantia* (LI *et al*. 2017) was used for CRISPR/Cas9 mutagenesis. *Exu-Cas9* was backcrossed to *LVPib12* each generation for stock maintenance.

### SPR mutant generation via CRISPR

*Spr^Δ235^*: We generated a mosquito line harboring a NHEJ-based deletion in exon 2 of *Spr* (AAEL019881), by CRISPR/Cas9-based editing according to the procedures in Kistler *et al*. (2015). Two sites for Cas9 cleavage targeting exon 2 were identified using CHOPCHOP (LABUN *et al*. 2016) and validated *in vivo*. Guide RNAs were synthesized *in vitro* using the MEGAScript kit (Thermo Fisher Scientific) from DNA generated from a template-free PCR consisting of a primer targeting the Cas9 cut site with a T7 promoter and a universal reverse primer (sequences listed in Table 1). After reaction purification with MEGAClear (Thermo Fisher Scientific) and size confirmation using a Bioanalyzer 2100 (Agilent Technologies), 40 ng/ul each of the two gRNAs together with 333 ng/ul Cas9 (PNA Bio) was injected into Thai strain embryos by the University of Maryland Insect Transformation Facility (https://www.ibbr.umd.edu/facilities/itf). After G_0_ females were backcrossed to Thai wild type males, blood fed and allowed to lay eggs, genomic DNA from the G_0_ mosquitoes was extracted using Puregene reagents (Qiagen) and PCR was used to detect deletions in the *Spr* gene (Table S1, Fig S1A). Eggs from deletion-positive females were hatched and back-crossed for five generations.

Heterozygous males and females were crossed, and progeny screened for genotype using genomic DNA extracted from a single leg (SMITH *et al*. 2018). Males and females homozygous for the *Spr* deletion were crossed together to generate a stable mutant line (*Spr^Δ235/Δ235^*). A wild type control line (*Spr^+/+^*) derived from a backcross of heterozygous *Spr^Δ235/+^* individuals was also maintained. *Spr^Δ235^* heterozygotes (*Spr^Δ235/+^*), derived from a cross between *Spr^Δ235^* homozygotes and this wild type line, served as an additional control.

*Spr^ECFP^*: We generated a disruptive insertion in exon 2 of *Spr* using CRISPR/Cas9-mediated homologous recombination. Guide RNAs were designed targeting exon 2 of *Spr* using CHOPCHOP (LABUN *et al*. 2016) and validated individually with *in vitro* cleavage assays using a PCR amplicon spanning the putative cut sites, each *in vitro* transcribed gRNA (MEGAscript, Invitrogen) and Cas9 protein (PNA bio). A single gRNA with validated cleavage activity was chosen for incorporation into a synthetic gBlock (IDT) that contained the *Ae*. *aegypti U6* (*AAEL017774*) promoter, a modified gRNA scaffold and a terminator (CHEN *et al*. 2021) for subsequent subcloning into the backbone of the following HDR donor plasmid. We first integrated homology arms into the base donor plasmid pSL1180-HR-PUbECFP (Addgene #47917) (MCMENIMAN *et al*. 2014). Homology arms 1425 bp (left) and 1797 bp (right) in length flanking the gRNA cut site were amplified with CloneAmp (Takara) from a consensus genomic DNA clone covering *Spr* exon 2 and flanking introns (primers listed in Table S2). We PCR-amplified the gBlock using primers with 5’ InFusion adaptors (Table S2). Next, we integrated the gBlock fragment with the U6 expression cassette into the backbone of this donor plasmid at the NdeI restriction site using InFusion cloning. This yielded a final construct (pMW001) containing a U6 expression cassette and HDR cassette targeting *Spr* exon 2.

We microinjected 300 ng/ul of pMW001 (endotoxin-free) into the posterior pole of *Exu-Cas9* pre-blastoderm stage embryos (*LVPib12* strain) using an Eppendorf FemtoJet 4X. Transformed G_1_ larvae with constitutive ECFP fluorescence were identified by screening for fluorescent bodies (PUb-ECFP) at the L3-L4 stage. Transgenic animals were then outcrossed to *LVPib12* for two generations before crossing the lines to generate a homozygous viable *Spr* mutant strain (*Spr^ECFP^*^/*ECFP*^). Insertion of the disruptive donor cassette into *Spr* exon 2 was confirmed by PCR amplification of genomic DNA using a three primer genotyping strategy (Table 2, Fig S1B) where one forward primer was centered on the CRISPR cut site so that it would only anneal to the wild type allele, one forward primer was placed in the polyubiquitin (PUb) sequence in the integrated cassette, and one reverse primer was nested in the right homology arm. This yielded a 342 bp amplicon for the wild type allele with an intact gRNA site, and a 524 bp amplicon for the mutant allele.

### Mating, fertility, and body size of the *Spr^Δ235^* allele in Thai background

Mating was examined by mating *en masse* two-to-three-day-old homozygous (*Spr^Δ235/Δ235^*) or heterozygous (*Spr^Δ235/+^*) females to three-to-four-day-old heterozygous males. Mating was performed in an 8 L container for 24 h in the absence of 10% sucrose. Mated females were dissected to confirm mating status by scoring the presence of sperm in the spermathecae. Eggs from both homozygous and heterozygous females mated to heterozygous males had similar hatch rates. Two biological replicates were performed from independently hatched cohorts. To determine relative body sizes of the mosquitoes, we used wing-length measurements as a proxy (NASCI 1990).

### Post-mating receptivity

#### *Spr^Δ235^* allele in Thai background

A single two-to-three-day-old homozygous (*Spr^Δ235/Δ235^*) or heterozygous (*Spr^Δ235/+^*) female was released into an 8 L container with 10 three-to-four-day-old heterozygous (*Spr^Δ235/+^*) males and observed closely. Once the female mated (maintained a copula for >8 sec), the mating pair was immediately collected and the female was placed in a 0.5 L cup. The mated male was excluded from additional matings and the mating arena was replenished with an additional virgin male. Two three-to-four-day-old males with dsRed-marked sperm were introduced into each 0.5 L cup either immediately or 24 h after the first female mating and then held together for 24 h. The lower reproductive tract of each female was then dissected and the spermathecae and bursa examined under a fluorescence microscope for the presence of red sperm along with non-fluorescent sperm, indicative of remating. As a control, individual virgin homozygous females or virgin heterozygous females with two dsRed-sperm males were placed together in 0.5 L cups for 24 h (n=10 for each genotype for each replicate) and the dissected reproductive tracts of females were examined to make sure the dsRed-sperm males could mate successfully with *Spr^Δ235^* mutant females. Two biological replicates from independently hatched cohorts were conducted. The genotype order of initial matings was reversed in replicate two to avoid bias.

#### *Spr^ECFP^* in LVPib12 background

To similarly test if *Spr^ECFP^* mutants were refractory to remating, a single four-to-five-day-old homozygous (*Spr^ECFP/ECFP^*) or heterozygous (*Spr^ECFP/+^)* female was released into a 355 ml insulated solo cup with 10 five-to-six-day-old wild type (*Spr^+/+^*) *LVPib12* males and observed closely. Once the female mated (maintained a copula for >8 sec), the mating pair was immediately collected. The male was removed and discarded, and the female was placed in a separate solo cup with other newly mated females. This mating scheme was then replicated until the target number of copulated females per genotype was obtained. After 1-1.5 h following their first mating, these females were introduced into a 20.3 x 20.3 x 20.3 cm cage (Bioquip, 1450A) with five-to-six-day-old males with dsRed-marked sperm (SMITH *et al*. 2007). The female to male ratio was 1:2 and they were held together for 24 h at which point males were removed by aspiration. The lower reproductive tract of each female was then dissected and the spermathecae examined under a fluorescence microscope for the presence of red sperm, indicative of remating.

Control matings were performed between homozygous (*Spr^ECFP/ECFP^*) females (n=32) and wild type (*Spr^+/+^*) *LVPib12* males as described above to estimate insemination rates after the first mating (31/32 mated). As a supplemental control to validate mating compatibility in this post-mating receptivity assay, homozygous (*Spr^ECFP/ECFP^*) females (n=20) were placed with two times the number dsRed-sperm males in a 20.3 x 20.3 x 20.3 cm cage (Bioquip, 1450A) for 24 h and the dissected reproductive tracts of females were examined for the presence of red sperm (20/20 mated). Finally, twenty virgin females of each genotype (*Spr^ECFP/ECFP^*and *Spr^ECFP/+^*) were dissected to confirm virginity (20/20 for each genotype were unmated).

### *Spr^ECFP^* Oviposition Assays

For oviposition assays with virgin or mated wild type (*Spr^+/+^*), heterozygous (*Spr^ECFP/+^*) or homozygous (*Spr^ECFP/ECFP^*) females, three-to-five day old females were blood fed to repletion using anesthetized Swiss-Webster mice (Johns Hopkins University Animal Care and Use Committee, Approval Number: MO21H373). For assays with mated females, all crosses were made between virgin females of the target genotype and two-day-old wild type virgin males. Crosses were established with a sex ratio of 1 female: 2 males in a 20.3 x 20.3 x 20.3 cm cage (Bioquip, 1450A) with constant access to 10% sucrose. These small group crosses were left to mate for 48h before blood feeding the females. 72 hours post blood feeding, females were placed in single oviposition vials, each containing a filter paper cone moistened with 4mL dH_2_O (WOHL AND MCMENIMAN 2023b). Females were allowed to lay eggs on the filter paper for 48 hours at which point they were removed from vials and eggs were counted. Egg papers from females who died before collection were not counted. Counts were analyzed statistically as described below.

#### Drosophila sex peptide injection

Synthetic sex peptide (3 pmol; CanPeptide) in 69 nl *Aedes* saline (HAYES 1953) was injected into the thorax of virgin two-to-five-day-old Thai background *Ae. aegypti* (n=18) and Canton S *D. melanogaster* females (n=30) with a Nanoject II (Drummond, Broomall, PA). Controls included injection with *Aedes* saline and a non-injected group. Males were introduced (ratio 1:1) 12 h after injection. Mating events were confirmed by immediate direct observation for *Drosophila*. The number of eggs laid by females in the treatment groups was recorded. *Ae. aegypti* females were examined for sperm presence or absence after being held with males for two days.

To independently test if *D. melanogaster* sex peptide (dSP) could induce oviposition behavior in the *Ae. aegypti LVPib12* genetic background, we injected synthetic dSP (Aapptec, sequence: WEWPWNRKOTKFOIOSONORDKWCRLNLGPAWGGRC with a disulfide bridge between the two cysteines) (CHEN *et al*. 1988) dissolved in PBS (Gibco, pH 7.2) into the thorax of virgin females. We injected a concentration series spanning the dynamic range of dSP concentrations known to elicit oviposition responses in *D. melanogaster*. Specifically, we performed 150 nl injections of 10 uM, 100 uM and 1 mM dSP which equates to 1.5 pmol, 15 pmol and 150 pmol of dSP respectively. First, three-to-five-day-old virgin females were blood fed using anesthetized Swiss-Webster mice, and 48 h later, synthetic dSP or *D. melanogaster* male accessory gland homogenate (dMAG), *Ae. aegypti* male abdominal tip (aeMAT) homogenate (preparation described in next section) and PBS controls were injected into the thorax with a Nanoject II (Drummond). Oviposition was assayed as described above.

### Accessory Gland Homogenate Injection Assays

We tested both the effects of injecting *Drosophila* and *Aedes* male accessory gland (MAG) homogenates on mating and egg laying:

To first evaluate the effect of *Drosophila* MAG (dMAG) on Thai *Ae. aegypti* mating behavior and oviposition, accessory glands were dissected from 38 six-day-old virgin *D. melanogaster* Canton S males in 38 μl *Aedes* saline (HAYES 1953). The tissues were ground, sonicated in a water bath for 15 sec, and then spun at 13,400 rpm for 15 minutes at 0°C. The supernatant was removed and virgin Thai *Ae. aegypti* females (three-to-six days old) were injected in the thorax with 0.25 μl of *D. melanogaster* homogenate (equivalent to 0.25 of an accessory gland). Females injected with *Aedes* saline were used as controls as well as females injected with *Ae. aegypti* male accessory gland extract (aeMAG); and females that were mated without injection. Females were divided into two cohorts. One group was tested for refractoriness to mating by placing females in individual 0.5 L cups with males (aged seven-to-nine days) 3 days after injection for 2 days (n=10). Two days later, females were removed and their spermathecae dissected to determine insemination status. The other group was tested for oviposition behavior. Females were blood fed on a human arm (M.E.H.) 4 days after injection. Unfed females were removed and offered blood the following day. Three to four days after feeding, fed females were placed in individual cups (0.5 L) for oviposition. Four days later, females were removed and the number of eggs for each female was counted. Live females with zero eggs were dissected to determine insemination status (only when females had been exposed to males) and for the presence of fully developed eggs in ovaries. Virgin females injected with saline and non-injected females mated with males from the same cohort were used as controls.

In the second independent assessment of the impact of *Drosophila* and *Aedes* accessory gland extracts on egg laying, *Drosophila* MAGs were prepared by dissecting accessory glands from 50 fly abdomens (Canton-S strain) into 50 ul of PBS (Gibco, pH 7.2) and homogenized with a pellet pestle motor (Kimble). This mixture was centrifuged at 14,000 x g for 30 minutes at 4°C. The supernatant was applied to a 0.22 um filter column (Millipore) and centrifuged at 14,000 x g for 10 minutes at 4°C. The liquid flow through (dMAG) was kept at 4°C and injected within one week of preparation. *Aedes* accessory gland homogenates were prepared using an identical protocol from three-to-five-day-old *LVPib12* virgin males that had been anesthetized on ice with the following modifications: The male abdominal tip (MAT) was dissected into a drop of PBS by grasping the genital claspers with forceps and pulling the terminal abdominal segment away from the rest of the abdomen. This isolated the last segment of the abdomen including the male accessory glands and other reproductive organs. MAT from 200 males were dissected into 200 uL PBS (Gibco, pH 7.2) on ice (HELINSKI *et al*. 2012). The liquid flow through (aeMAT) was kept at 4°C and injected within two weeks of preparation.

To test if SPR is required for the post-mating reduction in mating receptivity resulting from injection of *Ae. aegypti* MAG (aeMAG) homogenate, Thai wild type and *Spr^Δ235^* homozygous females were injected with aeMAG as previously described (AMARO *et al*. 2021). Modified PBS buffer alone (137 mM NaCl, 2.7 mM KCl, 10 mM Na_2_HPO_4_, 3 mM KH_2_PO_4_, 2 mM CaCl_2_ pH 7.0) served as a control. Two days after injection, females were blood fed on a human host (S.P.) for 8 minutes, fully-engorged females were separated individually into 0.5 L cups and 2 wild type Thai males were introduced 5 days post-blood feeding. After 2 days female spermathecae were dissected and scored for the presence or absence of sperm.

### Cell-based assays

Cell-based assays were performed as in Duvall *et al*. (2019). Briefly, HEK293T cells (Thermo Fisher Scientific) were maintained using standard protocols in a Thermo Scientific water jacketed carbon dioxide incubator. Cells were transiently transfected with 0.5 μg each of plasmid expressing GCaMP6s (CHEN *et al*. 2013), mouse Gqα15 (OFFERMANNS AND SIMON 1995), and SPR (AAEL019881) using Lipofectamine 2000 (Invitrogen). Transfected cells were seeded into 96 well plates (Greiner Bio-one) and incubated overnight in Fluorobrite DMEM media (Thermo Fisher Scientific) supplemented with Fetal Bovine Serum (Invitrogen) at 37°C and 5% carbon dioxide. Cells were directly imaged in 60μL Fluorobrite DMEM media (Thermo Fisher Scientific) using GFP-channel fluorescence of a BioTek Synergy Neo plate reader with liquid handling system and automated dispenser. Peptides were prepared at 3x concentration in reading buffer [Hank’s Balanced Salt Solution (GIBCO), 20mM HEPES (Sigma-Aldrich), pH 7.4]. Wells were imaged every 0.5 seconds for 3 minutes in kinetic mode. 30μL of peptide was added to each well after 15 seconds of baseline fluorescence recording. Normalized responses were calculated as (ΔF/F_0_)_experimental_ – (ΔF/F_0_)_no receptor control_. AstAR (AAEL006076) response to Ast1 was used as a positive control in each plate. *Ae. aegypti* peptides previously detected in Predel *et al*. (2010) and tested *in vitro* in Duvall *et al*. (2019) were synthesized by Bachem and maintained as lyophilized powders or 100% dimethyl sulfoxide (DMSO) (Sigma-Aldrich) concentrated stock solutions at -20°C. All peptides were tested at a 10 μM final concentration with < 1% DMSO final concentration. 11 – 14 replicates were performed/transfection and each experimental well included matched positive and negative (no receptor transfected) control wells in the same plate. Any well in which the matched positive control response was < 0 was excluded from analysis.

#### Data analysis

Power analysis using G*power (FAUL *et al*. 2007) was used to determine appropriate sample size prior to experiments based on preliminary data. Statistical analysis was performed using SPSS (IBM Statistics, version 25). A Levene’s test was first performed to determine homoscedasticity of male and female body size data, then wing lengths were compared with Kruskal-Wallis and Mann-Whitney tests. Replicate effects were analyzed and replicates were combined when appropriate. Remating was analyzed using a binary generalized linear mixed model to compare differences by time points and genotype (*Spr^Δ235/Δ235^*or heterozygous (*Spr^Δ235/+^*); to compare remating in the *Spr^ECFP^* mutants vs. wild type, a two sample proportions test was performed. Egg counts from oviposition assays were analyzed with Graphpad Prism 9.4.1 software with a one-way ANOVA followed by *post-hoc* t-tests between experimental groups. *P* values were adjusted for multiple comparisons with Dunnett’s correction when the comparisons were all against a control group; and adjusted with Tukey’s correction when comparisons were made between experimental groups. In cell-based assays fluorescence signal responses were analyzed with Graphpad Prism 9.3.1 software with one sample t and Wilcoxon Signed Rank test.

## Results

### Generation and characterization of *Aedes aegypti Spr* mutants

To test whether sex peptide receptor (SPR) is required for canonical post-mating receptivity and egg laying behaviors in *Ae. aegypti*, we generated two independent mutant alleles in the wild type *Spr* gene locus using CRISPR/Cas9 mutagenesis (Fig 1A).

**Figure 1:**
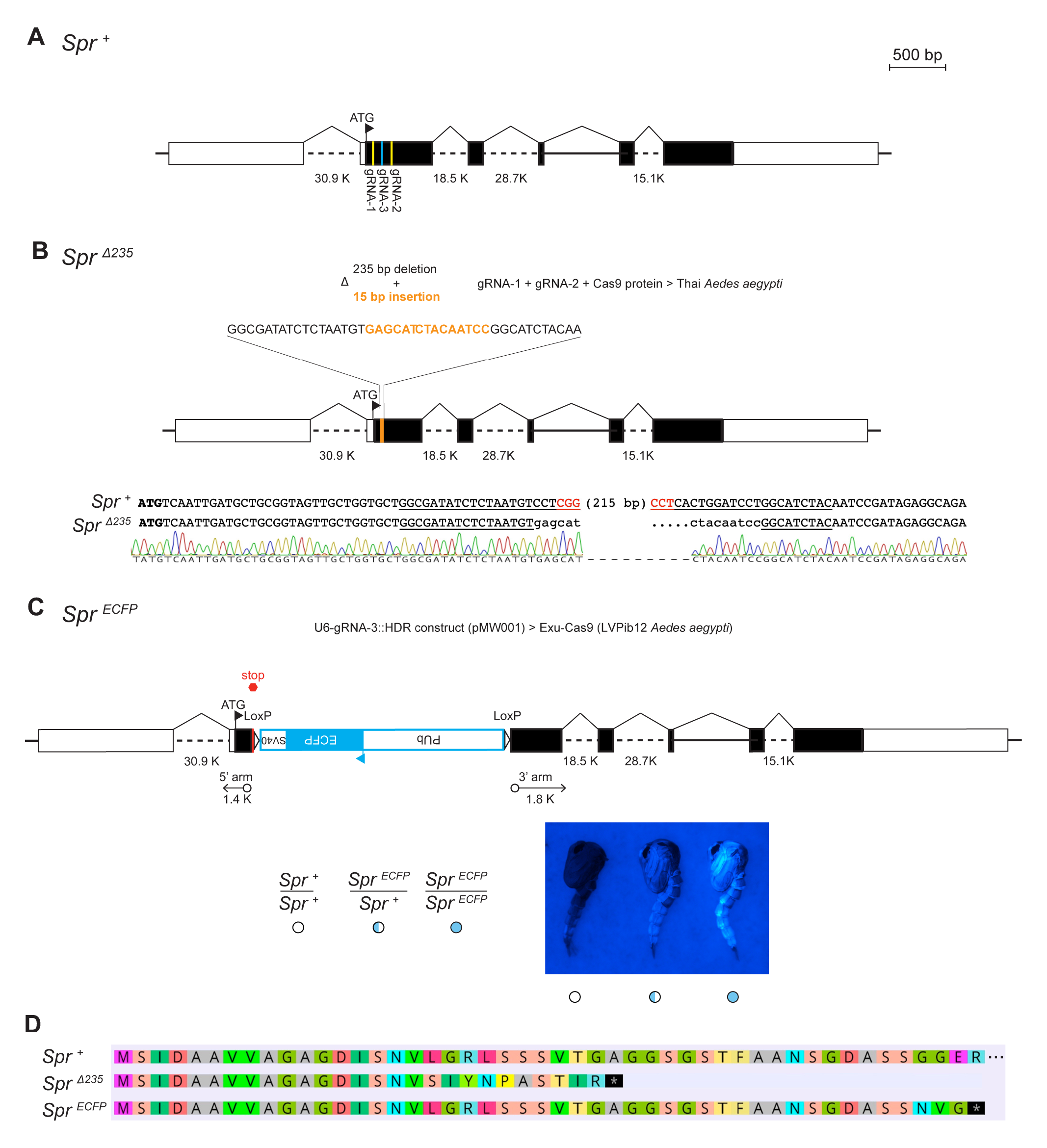
Generation of *Aedes aegypti* sex peptide receptor mutants. (A) Diagram of the wild type *Ae. aegypti* sex peptide receptor (*Spr^+^*) genomic locus. Boxes denote exons which are shaded with white for untranslated regions and black for the open reading frame. Guide RNA sites used for generating mutants shown as yellow and blue lines. (B) Diagram of the *Spr^Δ^*^235^ mutant allele generated by deletion of 235 bp and insertion of 15 bp using gRNA1 and gRNA2. Sanger trace (below) validates the altered sequence of this allele relative to wild type. Translational start site is bold, gRNA target site underlined, and PAM sequence highlighted in red. (C) Diagram of *Spr ^ECFP^* mutant allele generated by insertion of a disruptive ECFP expression cassette via homology directed repair. Representative images of wild type (open circle), heterozygous (half cyan circle) and homozygous (full cyan circle) *Spr^ECFP^*mutant pupae are shown below. (D) Predicted amino acid alignments of wild type and truncated *Spr^Δ^*^235^ and *Spr^ECFP^*mutant alleles. Premature stop codons are indicated by the black box with star.

We first generated a NHEJ allele (*Spr^Δ235^*) in the Thai *Ae. aegypti* genetic background by injecting two gRNAs targeting exon 2 of *Spr* (the first coding exon of this gene) along with Cas9 protein into pre-blastoderm stage embryos (Fig 1B). Subsequently using PCR genotyping (Fig S1A), we successfully recovered the *Spr^Δ235^* mutant allele consisting of a 235 bp deletion (bp 50-285 in the open reading frame) with an insertion of 15 bp in this exon.

We next generated an independent HDR allele (*Spr^ECFP^*) in the *LVPib12 Ae. Aegypti* genetic background by injecting a donor construct with a gRNA expression cassette in its backbone into pre-blastoderm stage embryos from the germline *Exu-Cas9 Ae. aegypti* strain (Fig 1C). In this allele, a 2617 bp cassette was inserted by homology-dependent repair into exon 2 of *Spr*. This disruptive insertion included a stop cassette and marks mutants visibly with enhanced cyan fluorescent protein (ECFP) (Fig 1C) and can be detected in the *Spr* locus by a custom genotyping assay using PCR (Fig S1B).

Predicted amino acid alignments indicate that relative to the wild type *Spr^+^* allele, both *Spr^Δ235^* and *Spr^ECFP^* mutant alleles are likely truncated before the first predicted transmembrane domain of SPR (Fig 1D). Specifically, the *Spr^Δ235^* allele has a frameshift mutation in the *Spr* coding sequence that yields a premature stop codon at amino acid 28, while precise insertion of a stop codon in *Spr^ECFP^* directly after amino acid 47 is similarly predicted to prematurely terminate translation.

### *Spr* knockout mutants are viable and fertile

We determined that both of the *Spr* mutant alleles that we generated (*Spr^Δ235^* and *Spr^ECFP^*) were homozygous viable. We have successfully maintained these homozygous mutant lines for more than six years and one year respectively since their establishment, using standard protocols.

Detailed morphometric and fitness characterization of the *Spr^Δ235^* allele, revealed that homozygous mutant males were smaller (average = 1.99 to 2.16 mm wing length) than wild type (*Spr^+/+^*) or heterozygous (*Spr^Δ235/+^*) males reared under same conditions (2.13 to 2.23 mm) across replicates and remating experiments; Z = -3.79, P<0.00 (wild type), Z = -4.05, P<0.00 (heterozygous). Mutant females were smaller in body size compared to wild type and *Spr^Δ235/+^* females in replicate 1: *Spr^Δ235/Δ235^* female = 2.53 ± 0.15 mm; *Spr^+/+^*female = 2.78 ± 0.20 mm (Z = -4.039, df <0.01), but not significantly different in Replicate 2: *Spr^Δ235/+^* female = 2.9 ± 0.06 mm; *Spr^Δ235/Δ235^*female = 2.88 ± 0.08; *Spr^+/+^* female = 2.91 ± 0.05. Mutant females mated successfully with males at the same percentage as controls, as demonstrated by the presence of sperm in their spermathecae after 24 h [96% *Spr^Δ235/Δ235^*(87/90) and 99% *Spr^Δ235/+^* (91/92) females mated with *Spr^Δ235/+^*males].

### SPR *in vitro* response screen

The smaller body size of our *Spr* mutants prompted us to investigate the nature of the ligand of SPR. Work from Kim *et al*. (2010) suggests that *Aedes* SPR can be potently activated by non-SP, non-MIP ligands found in whole mosquito extracts. We performed *in vitro* assays to profile *Ae. aegypti* SPR responses to various classes of neuropeptides. We expressed *Ae. aegypti* SPR cDNA in HEK293T cells co-transfected with a promiscuous G protein (OFFERMANNS AND SIMON 1995) and the genetically encoded calcium sensor GCaMP6s (CHEN *et al*. 2013). Activation of a given receptor is read out by calcium-induced increase in fluorescence of GCaMP6s. SPR responses were profiled to a 10μM dose of 50 known neuropeptides. Although modest responses were noted to several neuropeptides (Ast3, AT, sNPF1, sNPF2+4, TKRP2, TKRP3), these showed low efficacy compared to positive controls (AstAR activation by Ast1 peptide) (Fig S2). We did not observe activation by MIP peptides tested, nor by HP-I, a peptide known to play a role in short-term receptivity suppression (DUVALL *et al*. 2017).

We did not identify additional high efficacy agonists of SPR among our neuropeptide panel.

### *Spr* knockout females show normal changes in post-mating refractoriness

Mated *Ae. aegypti* females quickly become refractory to subsequent mating (CRAIG 1967; HELINSKI *et al*. 2012; DEGNER AND HARRINGTON 2016). While initial changes are induced by a seminal peptide HP-1 (DUVALL *et al*. 2017), this effect is temporary, acting only within one hour of mating. The nature of the molecule(s) that induce(s) lifelong refractoriness in these females is unknown.

We initially observed that *Spr^Δ235/Δ235^* homozygous mutant females are as likely to mate with a second male as are *Spr^Δ235/+^* control females, regardless of the time after the first mating (Fig 2A; p = 0.88 and p = 0.18 for tests immediately, and 24 h, after mating, respectively). To confirm this result, we injected male accessory gland homogenate, which is known to induce mating refractoriness (HELINSKI *et al*. 2012), from wild type Thai males into both *Spr^Δ235/Δ235^*and wild type Thai females and found no difference in mating receptivity when males were presented 5 days after injection [*Spr^Δ235/Δ235^*, 1.37% females remated (n = 74); wild type, 1.16% females remated (n = 86)].

**Figure 2:**
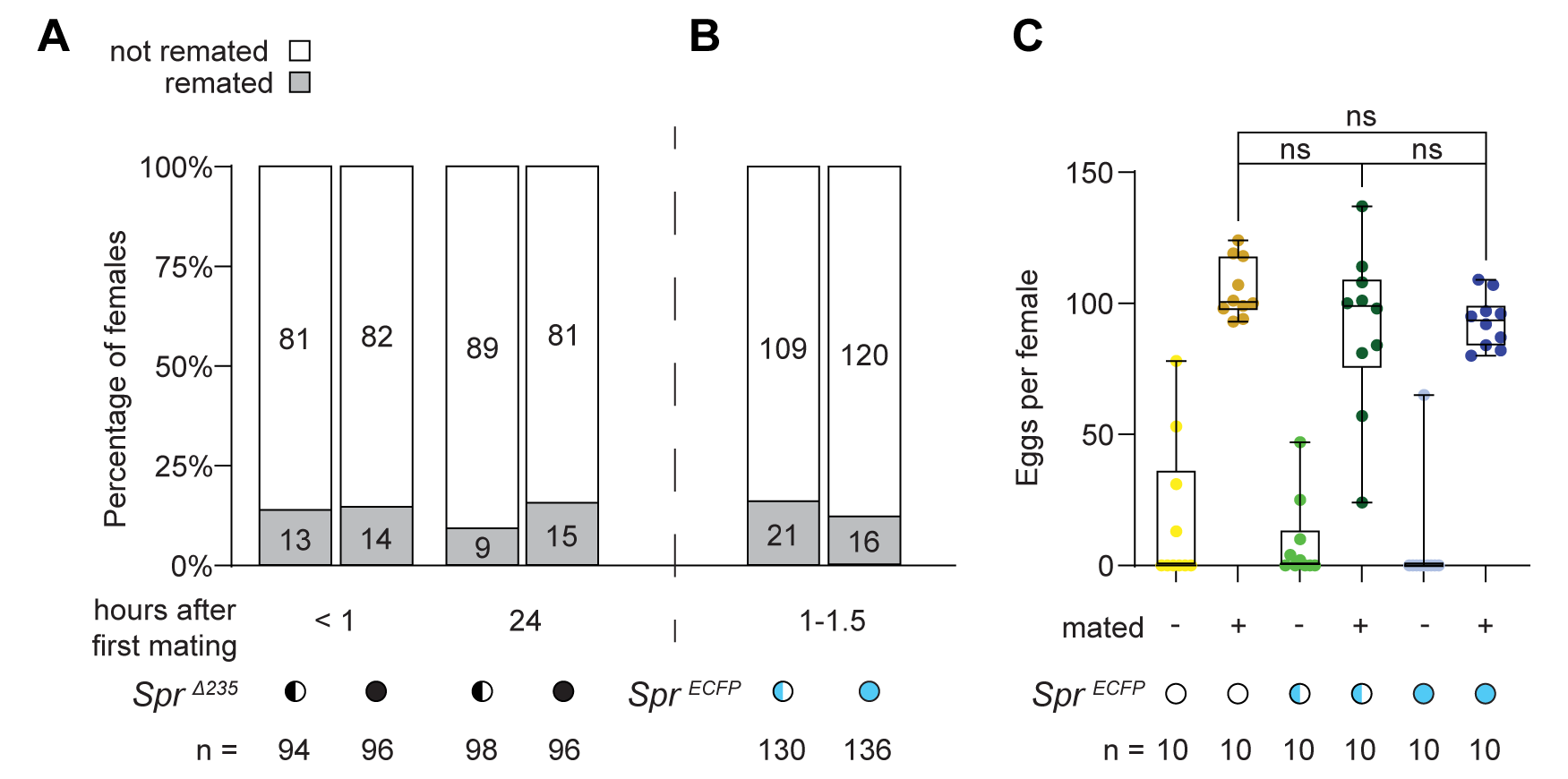
Sex peptide receptor is not required for refractoriness to remating over varied timescales or induction of egg laying in *Aedes aegypti*. (A) Percent of *Spr^Δ235^* heterozygous (half black circle) and homozygous mutant (full black circle) females that remated (gray shading) immediately (<1 hr) or 24 h after their first mating. Raw number of females in each remating category are denoted within each bar. Cumulative sample size (n) from two independent biological replicates indicated. (B) Percent of *Spr^ECFP^* heterozygous (half cyan circle) and homozygous (full cyan circle) mutant females that remated (gray shading) 1 to 1.5 h after their first mating. Raw number of females in each remating category are denoted within each bar. Sample size indicated below genotypes. (C) Eggs laid per female for wild type (open circle), *Spr^ECFP^* heterozygous (half cyan circle) and *Spr^ECFP^* heterozygous (full cyan circle) mutant genotypes after a bloodmeal that were either virgin (-) or mated (+). n.s. = p > .05 (Dunnett’s multiple comparisons test). Box plots show median, interquartile range, and maximum and minimum values.

Similarly, we observed in an independent post-mating receptivity assay, that *Spr^ECFP/ECFP^* homozygous mutant females are as likely to mate with a second male as are *Spr^EFCP/+^* control females (Fig 2B; Z = 0.88, p = 0.37). In this assay, females were tested for their propensity to remate within 1-1.5 h after initial mating and we similarly detected no significant differences in post-mating receptivity.

Thus, *Spr* does not seem to be necessary for induction of post-mating refractoriness, including short, medium or long-term refractoriness, in *Ae. aegypti* females.

### *Spr* is not required for egg laying in *Aedes aegypti*

Mating stimulates oviposition in gravid *Aedes aegypti* females (LANG 1956; JUDSON 1967), and this effect is mediated by unknown protein/s transferred to females in male accessory gland fluid (LEAHY AND CRAIG 1965; HISS AND FUCHS 1972). To test whether SPR is required for post-mating oviposition behavior in *Ae. aegypti*, we next tested whether egg laying was impacted using the *Spr^ECFP^* mutant allele.

We determined that gravid wild type *Spr^+/+^*, heterozygous *Spr^ECFP/+^* and homozygous *Spr^ECFP/ECFP^* mutant females laid very few eggs if they were not mated: 6.5 ± 6.5, 8.8 ± 4.9 and 17.5 ± 8.8 eggs, respectively (mean ± S.E.M., Fig 2C). We hypothesized that if SPR was required for post-mating oviposition behavior, homozygous *Spr^ECFP/ECFP^* females would not lay eggs even after mating. However, the number of eggs laid by mated homozygous *Spr* mutant females (90.4 ± 10.0 eggs) was not significantly different from that laid by heterozygous (92.9 ± 3.1 eggs, comparison: p = 1.000) or wild type females (mean: 105.3 ± 3.5, comparison: p = 0.774) (mean eggs ± S.E.M., Fig 2C).

We conclude that SPR is not required for post-mating oviposition in *Ae. aegypti*.

### Neither *Drosophila melanogaster* sex peptide nor male accessory gland homogenate induces mating refractoriness in *Aedes aegypti* females

We next tested whether injection of synthetic *D. melanogaster* SP (dSP) could induce mating refractoriness in *Ae. aegypti,* as demonstrated for *D. melanogaster* and *H. armigera* (CHEN *et al*. 1988; SCHMIDT *et al*. 1993; FAN *et al*. 1999; FAN *et al*. 2000).

We first confirmed activity of synthetic dSP by injecting 3 pmol (or saline) into 30 virgin *D. melanogaster* and assessing activity. Mating refractoriness assays revealed that only 6.6% of SP-injected *D. melanogaster* female flies mated, compared with 63% of those injected with saline. Similarly in egg laying assays, 46.7% of SP-injected *D. melanogaster* virgin female flies laid eggs compared to 0% of *D. melanogaster* females injected with saline. These data are all consistent with previous reports of the effects of dSP on mating refractoriness and egg laying in *D. melanogaster* (CHEN *et al*. 1988; SCHMIDT *et al*. 1993; LIU AND KUBLI 2003).

We then examined the effect of injecting synthetic dSP (3 pmol) or saline vehicle on mating refractoriness in the Thai *Ae. aegypti* strain using virgin females. In these assays, at 12 h post-injection, injected females were allowed to mate for 48 h and afterwards, females were dissected and examined for sperm in their spermathecae. We observed that sperm were present in all dSP-injected and all saline-injected females, indicating that dSP had no effect on *Ae. aegypti* mating receptivity in this context (Fig 3A).

**Figure 3:**
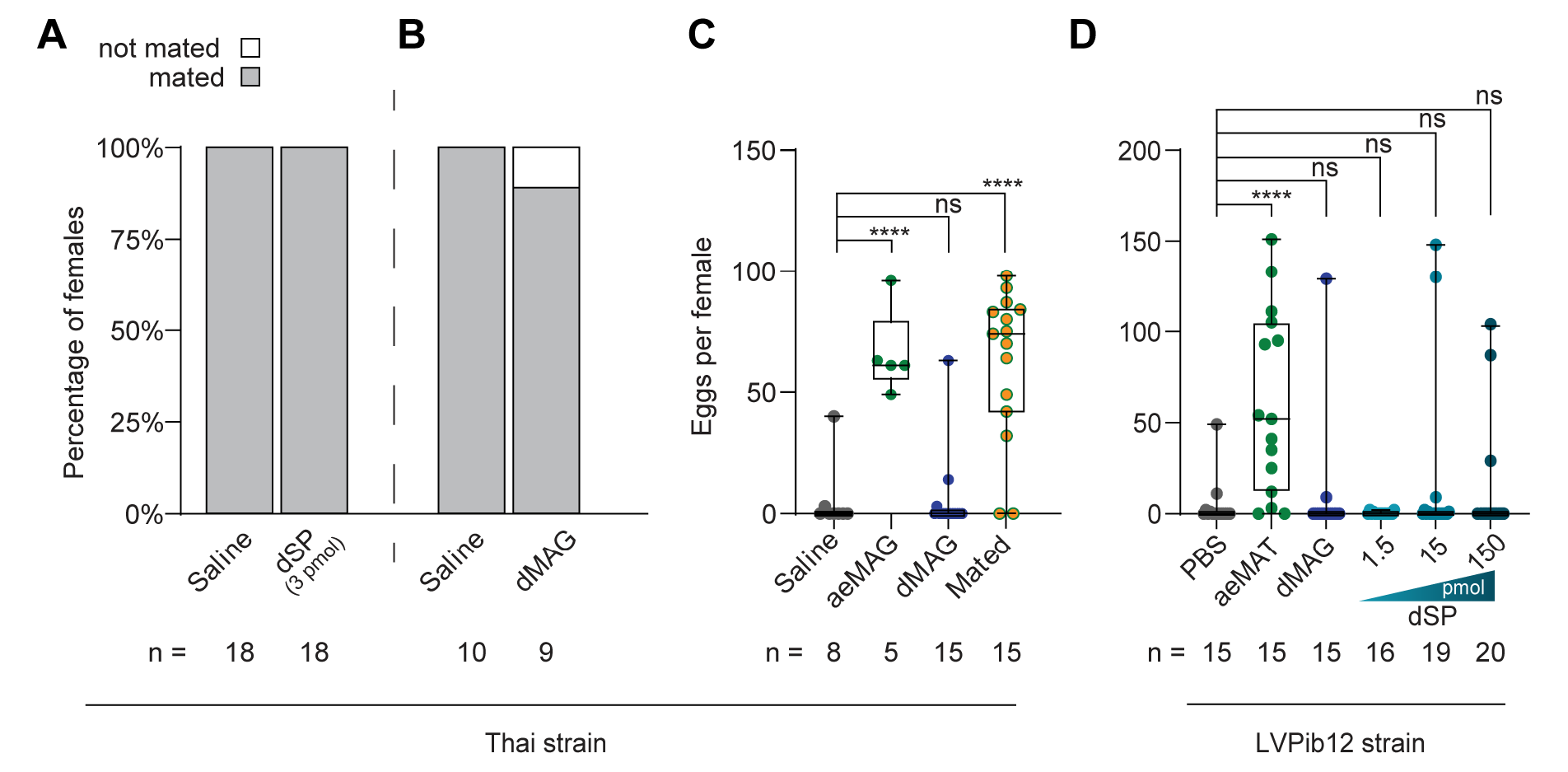
*Drosophila melanogaster* sex peptide and accessory gland extracts do not induce post mating behaviors in *Aedes aegypti* females. (A) Percentage of virgin *Ae. aegypti* females injected with saline vehicle or synthetic *Drosophila* sex peptide (dSP) that mated (gray shading). (B) Percentage of virgin *Ae. aegypti* females injected with saline vehicle or *Drosophila* male accessory gland (dMAG) homogenate that mated (gray shading). (C) Eggs laid per virgin gravid *Ae. aegypti* female after injection with saline vehicle or male accessory gland homogenate from *Aedes aegypti* (aeMAG) or *Drosophila melanogaster* (dMAG), or normal mating. n.s. = p > .05, **** p < .0001 (Dunnett’s multiple comparisons test). (D) Eggs laid per virgin gravid *Ae. aegypti* female after injection with saline vehicle (PBS), male abdominal tip homogenate from *Aedes aegypti* (aeMAT), dMAG, or different dosages of dSP. n.s. = p > .05, **** p < .0001 (Dunnett’s multiple comparisons test). Box plots show median, interquartile range, and maximum and minimum values.

Consistent with our results with synthetic SP, when 0.25 male *D. melanogaster* male accessory gland (dMAG) equivalents were injected into virgin Thai *Ae. aegypti* females (0.25 ul homogenate), we also found no effect of dMAG on mating receptivity (Fig 3B). We conclude that intrathoracic injection of synthetic dSP as well as homogenate from dMAG, the source of endogenous SP, is insufficient to induce mating refractoriness in *Ae. aegypti*.

### Neither *Drosophila melanogaster* sex peptide nor male accessory gland homogenate induces egg laying in *Aedes aegypti* females

Intrathoracic injection of *Ae. aegypti* male accessory gland homogenate has previously been shown to be sufficient to stimulate oviposition behavior in virgin gravid *Ae. aegypti* females (LEAHY AND CRAIG 1965; JUDSON 1967; HISS AND FUCHS 1972). Independently, our two groups next assessed whether intrathoracic injection of *D. melanogaster* homogenates putatively representing the complement of seminal fluid proteins from this species, inclusive of dSP, could similarly act to induce oviposition in virgin *Ae. aegypti* females.

In our first assay series in the Thai *Ae. aegypti* strain, we confirmed that injection of *Ae. aegypti* accessory gland homogenate (aeMAG) triggers oviposition from virgin gravid females (66.0 ± 8.0), approaching levels of egg laying observed from mated females (62.1 ± 8.1) (mean eggs ± S.E.M, Fig 3C). In contrast, virgin gravid females injected with dMAG laid very few eggs (5.3 ± 4.2) not significantly different from saline injected controls (5.4 ± 5.0).

Our second assay series in the *LVPib12* strain similarly revealed a crude homogenate prepared from the terminal abdominal segment of virgin males (*Ae. aegypti* male abdominal tip: aeMAT) containing the accessory glands and all other male reproductive glands was sufficient to induce oviposition behavior when injected into the thorax of virgin gravid females. Females injected with only PBS solvent laid on average 4.2 ± 3.3 eggs whereas those injected with aeMAT laid on average 60.7 ± 13.0 eggs (mean eggs ± S.E.M. aeMAT vs PBS control, p < 0.0001, Fig 3D). In contrast, dMAG did not stimulate egg laying from virgin gravid *Ae. aegypti* females (9.2 ± 8.6 eggs), yielding a similar level of oviposition to those injected with the PBS solvent control (mean eggs ± S.E.M., dMAG vs PBS control: p = 0.992, Fig 3D).

Finally, to probe whether *Drosophila* sex peptide (dSP) was capable of stimulating oviposition behavior in *Ae. aegypti*, we injected synthetic *D. melanogaster* sex peptide (dSP) over a concentration series ranging from 1.5 pmol to 150 pmol (Fig 3D) into virgin gravid females from the *LVPib12* strain. We determined that dSP injection did not have an effect on egg-laying in this context. The number of eggs (mean eggs ± S.E.M.) laid by females injected with 1.5 pmol dSP (0.3 ± 0.2, vs PBS control: p = 0.997), 15 pmol dSP (15.3 ± 10.0, vs PBS control: p = 0.787) and 150 pmol dSP (11.0 ± 6.7, vs PBS control: p = 0.961) did not differ significantly relative to the PBS solvent control.

We conclude that intrathoracic injection of dMAG as well as synthetic dSP are insufficient to stimulate oviposition behavior in virgin gravid *Ae. aegypti* females. However, aeMAT and aeMAG is sufficient to stimulate oviposition behavior within this context.

## Discussion

Elucidating the molecular pathways involved in establishing post-mating responses in *Ae. aegypti* from both males and females is important to our understanding of their reproductive biology and has implications for strategies for vector control. Here, we explored the role of SPR, a critical female receptor in some other insects, including the dipteran *D. melanogaster,* in mediating multiple post-mating responses. The primary *D. melanogaster* SPR ligand responsible for controlling reproductive outcomes is SP, a seminal peptide that is not found in insects outside *Drosophila* genus (TSUDA AND AIGAKI 2016; MCGEARY AND FINDLAY 2020). Interestingly, injecting *D. melanogaster* SP into unmated female *H. armigera* moths reduces egg laying, suppresses pheromone synthesis, and reduces calling behaviors (FAN *et al*. 1999; FAN *et al*. 2000; HANIN *et al*. 2012), thus mimicking post-mating responses analogous to those induced by SP in *Drosophila*. However, roles of SPR, and effects of SP were not consistent across insects. For example, injection of SP^21-36^ into the tarnished plant bug *Lygus herperus* had no effect on mating receptivity and although this region of SP binds *in vitro* to SPR from *D. melanogaster* and *H. armigera* it was unable to bind to SPR from *L. herperus* (HULL AND BRENT 2014). Although *Ae. Aegypti* oviposition was reported to be induced by surgical implantation of whole *D. melanogaster* accessory glands (LEAHY 1967) or intrathoracic injection of whole *Drosophila* male body lysates in the thorax of blood fed, virgin females (HISS AND FUCHS 1972), it is not known what male molecule(s) had this effect nor what receptor they bound to in the female. It should be noted that the stimulation of oviposition with intrathoracic injection of *Drosophila* male accessory gland homogenate into gravid virgin *Ae. aegypti* females was not observed when independently tested by both groups. In contrast, we found that *Ae. aegypti* accessory gland homogenate preparations were fully sufficient to evoke the post-mating receptivity and egg laying phenotypes that we characterized in this study consistent with previous reports (HISS AND FUCHS 1972; HELINSKI *et al*. 2012).

Genes corresponding to a SP homolog are not readily identified in insects outside *Drosophila*, but sequences homologous to SPR can be identified in various insect groups, including *Ae. aegypti*.; the mosquito SPR even binds Drosophila SP *in vitro*, albeit weakly (KIM *et al*. 2010; LEE *et al*. 2020). Given the presence of SPR in *Ae. aegypti*, the parallels between post-mating responses in *Ae. aegypti* and *D. melanogaster*, and the importance of SP in inducing post-mating response in *H. armigera,* we tested here whether SPR is required for post-mating responses in *Ae. aegypti*. We generated two independent SPR null mutations by CRISPR/Cas9 gene editing and compared mutant and control females for a variety of post-mating responses.

First, we tested whether SPR is needed for the post-mating drop in female receptivity. Female *Ae. aegypti* that have mated are refractory to subsequent mating (CRAIG 1967; HELINSKI *et al*. 2012); ∼25% do not remate immediately after mating, and all the mosquitoes eventually establish strong refractoriness 16-20 h after the initial mating (DEGNER AND HARRINGTON 2016). In both of the *Spr* mutant alleles (*Spr^Δ235^* and *Spr^ECFP^*) that we characterized, functional knockout of SPR does not affect the establishment of remating refractoriness, regardless of the interval between mating opportunities.

Stimulation of egg development and oviposition is another post-mating response by *Ae. aegypti* females (LANG 1956; JUDSON 1967). *Spr^ECFP^* knockout females oviposited a similar number of eggs as heterozygous females after mating and blood feeding, demonstrating that SPR is not required for this post-mating process. Knockdown or knockout of SPR in various insect systems have noted a reduced number of eggs laid (LI *et al*. 2014; ZHENG *et al*. 2015; GREGORIOU AND MATHIOPOULOS 2020; LIU *et al*. 2021), although the degree of reduction varies and it often is not a complete lack of egg laying. Consistent with the lack of effect of SPR knockout on mating receptivity and on oviposition, no phenotypes were observed when two commercial sources of synthetic dSP (at a biologically-effective dose for *D. melanogaster*) were injected independently into unmated *Ae. aegypti* females. The lack of any observable impact of intrathoracic microinjection of synthetic sex peptide on *Ae. aegypti* egg laying thus mirrored that of injecting dMAG homogenate which had the effect as described above.

SPR was reported to bind myoinhibitory peptides (MIPs), often at stronger affinities than SP, leading to the hypothesis that SPR is an ancestral MIP receptor (KIM *et al*. 2010; POELS *et al*. 2010). MIPs in insects are involved in control of hindgut and oviduct muscle contraction (SCHOOFS *et al*. 1991; BLACKBURN *et al*. 1995; BLACKBURN *et al*. 2001; PALUZZI *et al*. 2015; LUBAWY *et al*. 2020), modulating nutritional preferences (HUSSAIN *et al*. 2016; MIN *et al*. 2016), maintaining sleep states in *D. melanogaster* (OH *et al*. 2014), as well as involvement in ecdysis (DAVIS *et al*. 2003; KIM *et al*. 2006a; KIM *et al*. 2006b; SANTOS *et al*. 2007). MIPs have been shown to have allostatic activity by inhibiting JH synthesis in the cricket *Gryllus bimaculatus* (LORENZ *et al*. 1995) and Brown-winged green bug *Plautia stali* (MATSUMOTO *et al*. 2017). The *B. mori* peptide PTSP inhibits ecdysone synthesis from the prothorasic gland (HUA *et al*. 1999). There are five identified MIPs in *Ae. aegypti* (PREDEL *et al*. 2010; SIJU *et al*. 2014) with positive staining in the CNS with an anti-MIP antibody (KIM *et al*. 2010), though intriguingly a non-MIP ligand for *Ae. aegypti* SPR was partially purified (KIM *et al*. 2010). In our *in vitro* assay we found that *Ae. aegypti* SPR showed modest responses to Ast3, AT, sNPF1, sNPF 2+4, TKRP2, TKRP3 but we did not observe significant responses to HP-I, a peptide known to be found in male accessory gland and transferred to females during mating (NACCARATI *et al*. 2012; DUVALL *et al*. 2017) nor did we identify novel high efficacy ligands for SPR among our cohort of 50 peptides. Unlike Kim *et al*. (2010), we did not observe responses from MIP peptides tested in this assay (MIP 1, 4, and 5), although we note that our assay differs in cell type and sensor, and that although a 10μM dose of peptide has been used successfully to identify ligand/receptor interactions (DUVALL *et al*. 2019) we acknowledge that a single dose of ligand may not be sufficient to identify all potential SPR agonists. Given no strong phenotype in the SPR knockout mosquitoes characterized in the assays presented here, it is still possible that there could be a subtle effect with other known functions of MIPs. It is also possible there are pathways where ligands other than MIPs bind SPR that were not included in our peptide panel or belong to a different class of ligands.

As highlighted above, SPR disruptions in different insects can have varying effects on reproductive traits. Our contribution characterizing the effects of lack of SPR in *Ae. aegypti* helps contribute to the evolutionary understanding of this key receptor. The SPR is flexible to evolve binding to similar ligands for different functions, some of which have been used in reproduction. Unknown signaling pathways operating during transfer of *Ae. aegypti* seminal fluid thus likely underlie long-term refractoriness to remating and induction of egg laying in this important disease vector.

### Data availability statement

Strains and plasmids are available upon request. The authors affirm that all data necessary for confirming the conclusions of the article are present within the article, figures, and tables.

## Acknowledgements

We thank NIH grant R01-AI095491 to LCH and MFW, NIH grant R21AI176101 to CJM, and NIH grant R35-GM137888 and a Beckman Young Investigator award to LBD for supporting this research. MPW was supported by postdoctoral fellowships from the NIH (T32AI007417) and a 2022 L’Oreal USA For Women in Science Fellowship. ASP was supported by an NSF GRFP award. We thank Dr. Ben Matthews for advice on CRISPR strategies for generation of the *Spr^Δ235^* mutant allele at Cornell and assistance with sequence interpretation, the University of Maryland Insect Transformation Facility for injecting the constructs, Dr. Erika Mudrak of the Cornell Statistical Consulting Unit for assistance with statistical analysis, Dr. Nilay Yapici for valuable discussions, and Dr. Garrett League, Lindsay Baxter, Elizabeth Martin, Jake Angelico, Natalie Bailey, Brady Dolan, Nicole Blattman, and Sean Lee for mosquito rearing and data collection assistance at Cornell. CJM thanks Bloomberg Philanthropies and Johns Hopkins Malaria Research Institute for generous supplemental funding and supporting infrastructure.

---

**Table S1.**
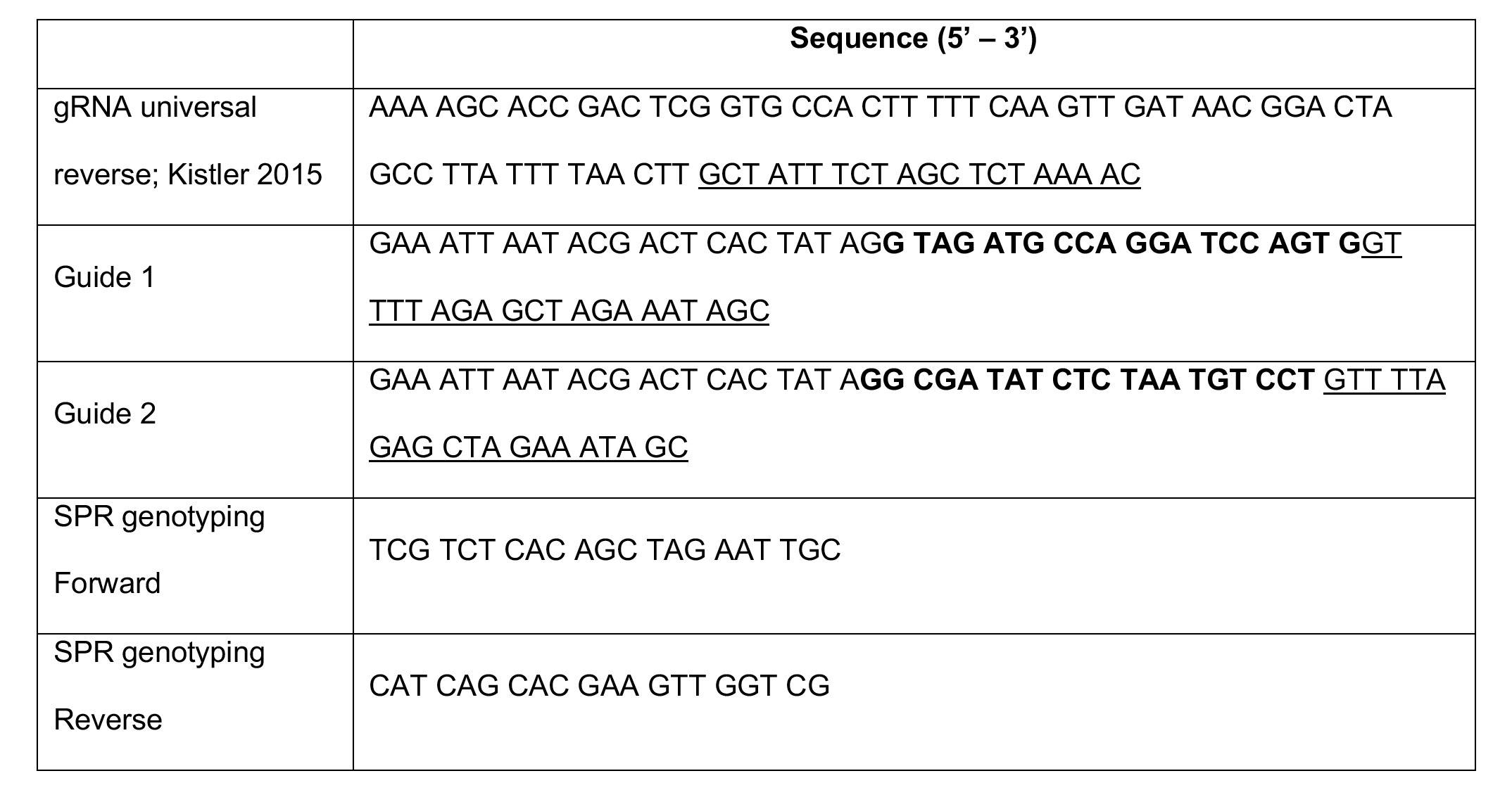
Primers for generation and verification of the *Spr^Δ235^* NHEJ allele. Underlined sequences in guide primers are complementary to the universal primer for use in template-free PCR to generate the DNA template for guide RNA synthesis. Bolded bases indicate the targeted *Spr* gene sequence.

**Table S2.**
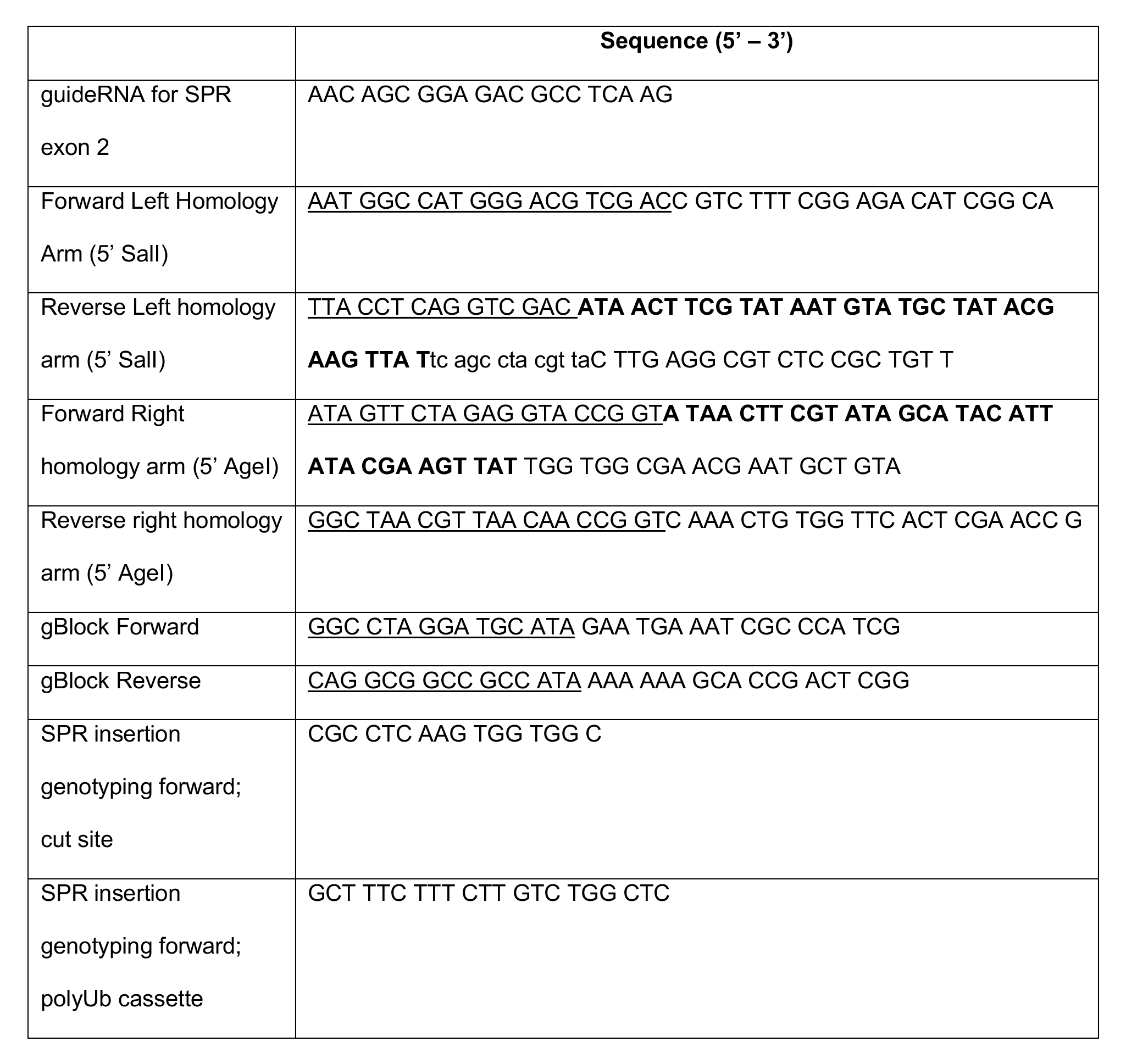
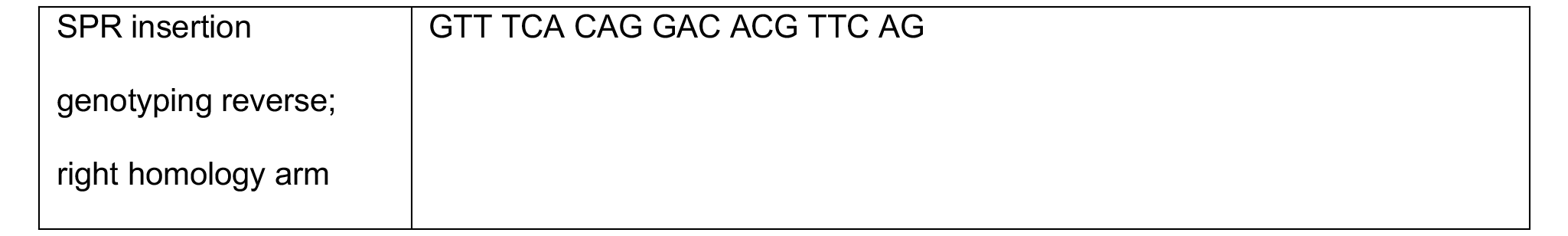
Primers for generation and verification of the *Spr^ECFP^* HDR allele. Underlined sequences indicate adapters for InFusion cloning, bold sequence indicates *LoxP* sites incorporated into the cassette to facilitate potential *Cre-LoxP* mediated excision of the integrated marker cassette, and lowercase letters indicate a triple stop cassette sequence.

**Figure S1:**
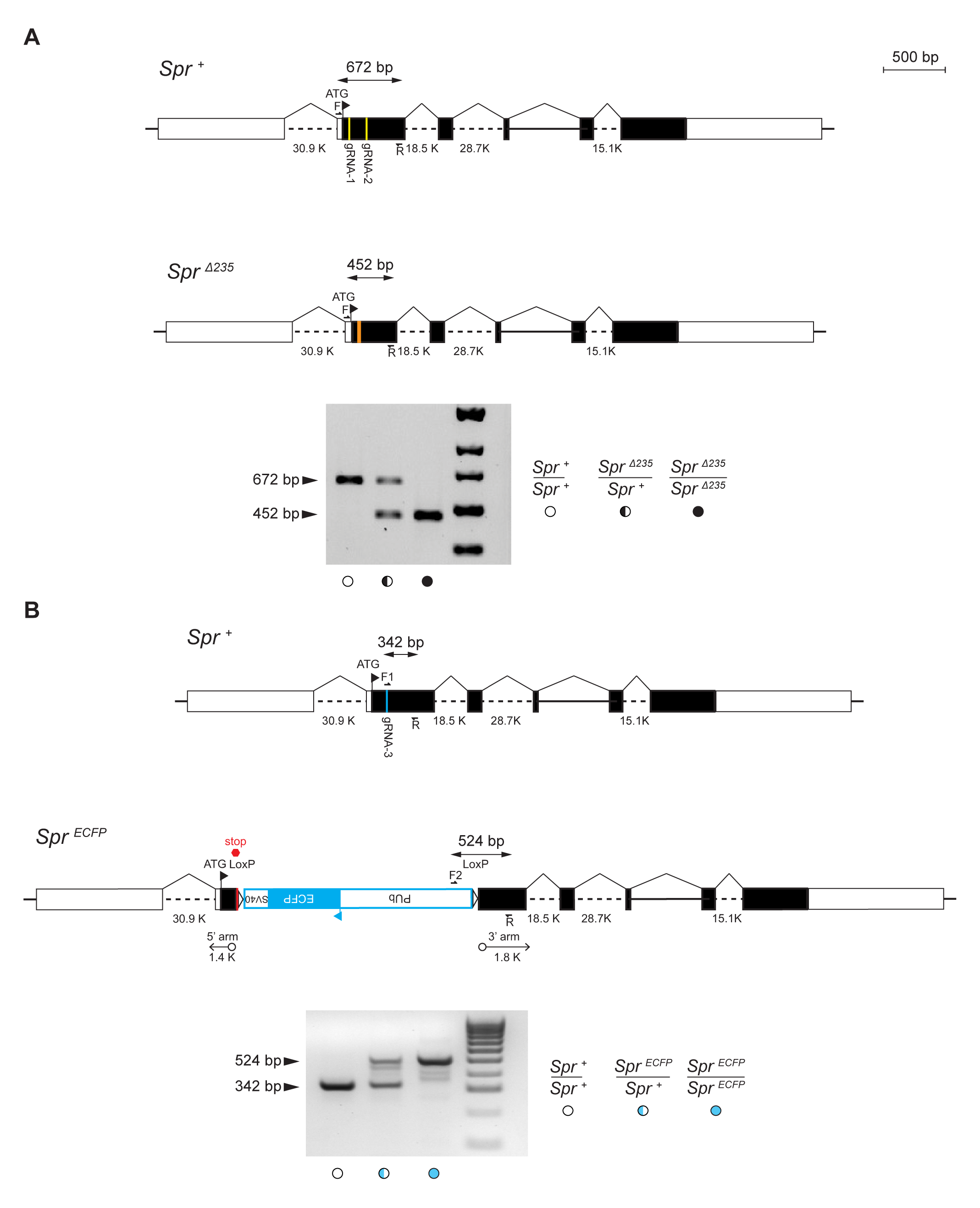
PCR genotyping assays for *Aedes aegypti* sex peptide receptor mutant alleles. (A) Genotyping assay for the *Spr^Δ235^*NHEJ allele: a two primer PCR assay with forward (F) and reverse (R) primers anchored in Exon 2 yields a 672 bp amplicon for the wild type *Spr^+^*allele and a 452 bp amplicon for the *Spr^Δ235^* mutant allele. (B) Genotyping assay for the *Spr^ECFP^* HDR allele: a three primer PCR assay was used where one forward primer (F1) was centered on the CRISPR cut site in Exon 2 so that it would only anneal to the wild type allele, one forward primer (F2) was placed in the polyubiquitin (PUb) sequence in the integrated cassette; and one common reverse primer (R) was nested in the right homology arm in Exon 2. This yields a 342 bp amplicon for the wild type *Spr^+^* allele with the intact gRNA site (amplified by F1/R), and a 524 bp amplicon for the *Spr^ECFP^* mutant allele (amplified by F2/R).

**Figure S2:**
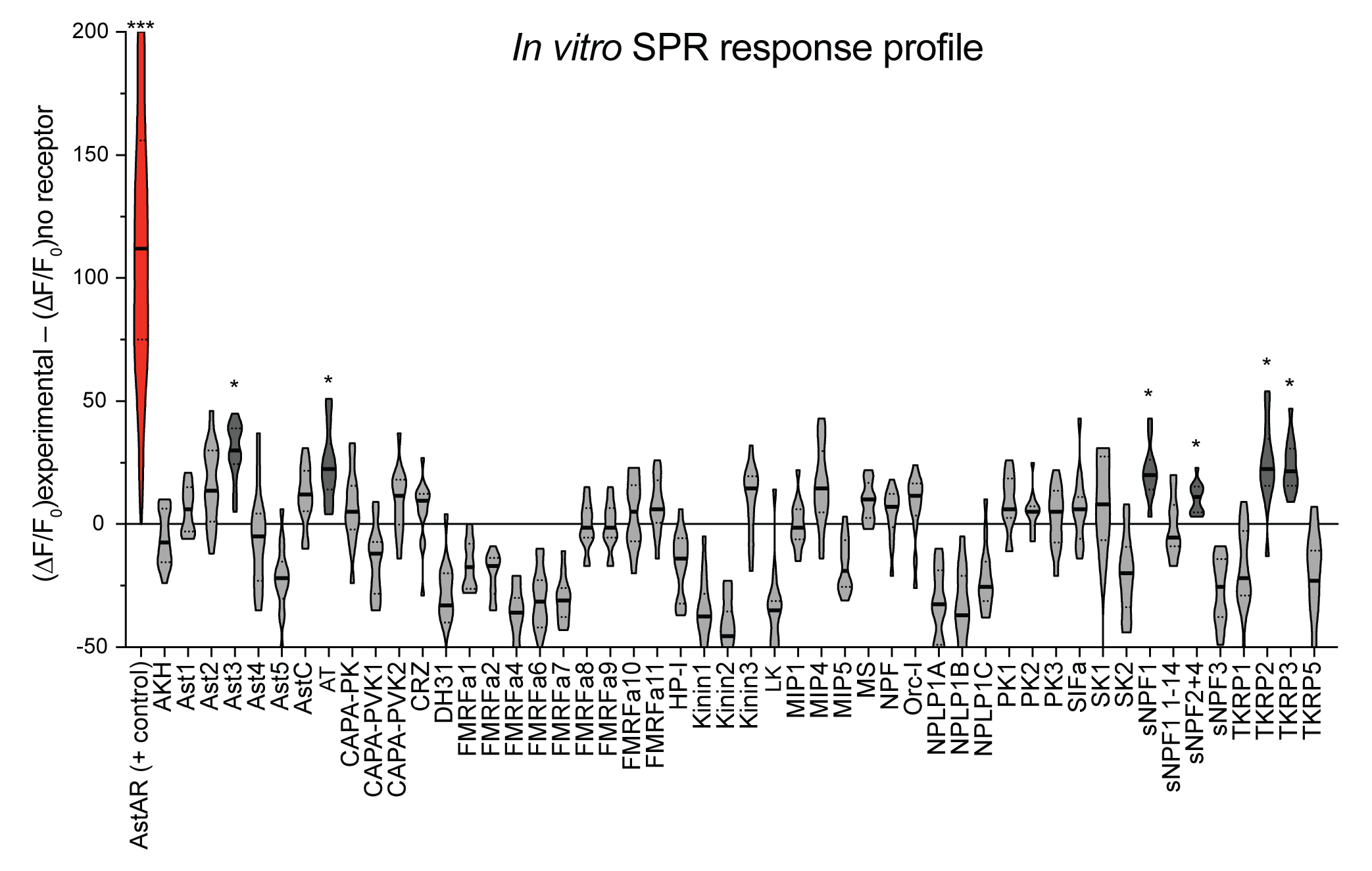
*In vitro Aedes aegypti* SPR responses to neuropeptide panel. *In vitro* response profile of *Ae. aegypti* sex peptide receptor (AAEL019881) to a panel of 50 neuropeptides. Fluorescence signal was calculated as (ΔF/F_0_)_experimental_ – (ΔF/F_0_)_no receptor transfected control._ AstAR (AAEL006076) response to Ast1 was used as a positive control in each plate. (n = 11 – 14 replicates/transfection). * p < 0.001, *** p < 0.0001 (one sample t and Wilcoxon Signed Rank test, hypothetical value = 0).

## References

Alfonso-Parra, C., Y. H. Ahmed-Braimah, E. C. Degner, F. W. Avila, S. M. Villarreal et al., 2016 Mating-Induced Transcriptome Changes in the Reproductive Tract of Female *Aedes aegypti*. PLoS Negl Trop Dis 10: e0004451.

Amaro, I. A., Y. H. Ahmed-Braimah, G. P. League, S. A. Pitcher, F. W. Avila et al., 2021 Seminal fluid proteins induce transcriptome changes in the *Aedes aegypti* female lower reproductive tract. BMC Genomics 22: 896.

Avila, F. W., A. L. Mattei and M. F. Wolfner, 2015 Sex peptide receptor is required for the release of stored sperm by mated *Drosophila melanogaster* females. J Insect Physiol 76: 1–6.

Avila, F. W., L. K. Sirot, B. A. LaFlamme, C. D. Rubinstein and M. F. Wolfner, 2011 Insect seminal fluid proteins: identification and function. Annu Rev Entomol 56: 21–40.

Blackburn, M. B., H. Jaffe, J. Kochansky and A. K. Raina, 2001 Identification of four additional myoinhibitory peptides (MIPs) from the ventral nerve cord of *Manduca sexta*. Arch Insect Biochem Physiol 48: 121–128.

Blackburn, M. B., R. M. Wagner, J. P. Kochansky, D. J. Harrison, P. Thomas-Laemont et al., 1995 The identification of two myoinhibitory peptides, with sequence similarities to the galanins, isolated from the ventral nerve cord of *Manduca sexta*. Regul Pept 57: 213–219.

Cator, L. J., C. A. S. Wyer and L. C. Harrington, 2021 Mosquito Sexual Selection and Reproductive Control Programs. Trends Parasitol 37: 330–339.

Chapman, T., J. Bangham, G. Vinti, B. Seifried, O. Lung et al., 2003 The sex peptide of *Drosophila melanogaster*: female post-mating responses analyzed by using RNA interference. Proc Natl Acad Sci U S A 100: 9923–9928.

Chen, H., H. Sun, J. Xie, Z. Yao, W. Zheng et al., 2023 CRISPR/Cas9-induced Mutation of Sex Peptide Receptor Gene Bdspr Affects Ovary, Egg Laying, and Female Fecundity in *Bactrocera dorsalis (Hendel)* (Diptera: Tephritidae). J Insect Sci 23.

Chen, J., J. Luo, Y. Wang, A. S. Gurav, M. Li et al., 2021 Suppression of female fertility in *Aedes aegypti* with a CRISPR-targeted male-sterile mutation. Proc Natl Acad Sci U S A 118.

Chen, P. S., E. Stumm-Zollinger, T. Aigaki, J. Balmer, M. Bienz et al., 1988 A male accessory gland peptide that regulates reproductive behavior of female *D. melanogaster*. Cell 54: 291–298.

Chen, T.-W., T. J. Wardill, Y. Sun, S. R. Pulver, S. L. Renninger et al., 2013 Ultrasensitive fluorescent proteins for imaging neuronal activity. Nature 499: 295–300.

Civetta, A., and R. S. Singh, 1998 Sex-related genes, directional sexual selection, and speciation. Mol Biol Evol 15: 901–909.

Craig, G. B., Jr., 1967 Mosquitoes: female monogamy induced by male accessory gland substance. Science 156: 1499–1501.

Davis, N. T., M. B. Blackburn, E. G. Golubeva and J. G. Hildebrand, 2003 Localization of myoinhibitory peptide immunoreactivity in *Manduca sexta* and *Bombyx mori*, with indications that the peptide has a role in molting and ecdysis. J Exp Biol 206: 1449–1460.

Degner, E. C., and L. C. Harrington, 2016 Polyandry Depends on Postmating Time Interval in the Dengue Vector *Aedes aegypti*. Am J Trop Med Hyg 94: 780–785.

Duvall, L. B., N. S. Basrur, H. Molina, C. J. McMeniman and L. B. Vosshall, 2017 A Peptide Signaling System that Rapidly Enforces Paternity in the *Aedes aegypti* Mosquito. Curr Biol 27: 3734–3742 e3735.

Duvall, L. B., L. Ramos-Espiritu, K. E. Barsoum, J. F. Glickman and L. B. Vosshall, 2019 Small-Molecule Agonists of *Ae. aegypti* Neuropeptide Y Receptor Block Mosquito Biting. Cell 176: 687–701 e685.

Fan, Y., A. Rafaeli, C. Gileadi, E. Kubli and S. W. Applebaum, 1999 *Drosophila melanogaster* sex peptide stimulates juvenile hormone synthesis and depresses sex pheromone production in *Helicoverpa armigera*. J Insect Physiol 45: 127–133.

Fan, Y., A. Rafaeli, P. Moshitzky, E. Kubli, Y. Choffat et al., 2000 Common functional elements of *Drosophila melanogaster* seminal peptides involved in reproduction of *Drosophila melanogaster* and *Helicoverpa armigera* females. Insect Biochem Mol Biol 30: 805–812.

Faul, F., E. Erdfelder, A. G. Lang and A. Buchner, 2007 G*Power 3: a flexible statistical power analysis program for the social, behavioral, and biomedical sciences. Behav Res Methods 39: 175–191.

Feng, K., M. T. Palfreyman, M. Hasemeyer, A. Talsma and B. J. Dickson, 2014 Ascending SAG neurons control sexual receptivity of *Drosophila* females. Neuron 83: 135–148.

Garbe, D. S., A. S. Vigderman, E. Moscato, A. E. Dove, C. G. Vecsey et al., 2016 Changes in Female *Drosophila* Sleep following Mating Are Mediated by SPSN-SAG Neurons. J Biol Rhythms 31: 551–567.

Gregoriou, M. E., and K. D. Mathiopoulos, 2020 Knocking down the sex peptide receptor by dsRNA feeding results in reduced oviposition rate in olive fruit flies. Arch Insect Biochem Physiol 104: e21665.

Haerty, W., S. Jagadeeshan, R. J. Kulathinal, A. Wong, K. Ravi Ram et al., 2007 Evolution in the fast lane: rapidly evolving sex-related genes in *Drosophila*. Genetics 177: 1321–1335.

Hanin, O., A. Azrielli, S. W. Applebaum and A. Rafaeli, 2012 Functional impact of silencing the *Helicoverpa armigera* sex-peptide receptor on female reproductive behaviour. Insect Mol Biol 21: 161–167.

Hasemeyer, M., N. Yapici, U. Heberlein and B. J. Dickson, 2009 Sensory neurons in the *Drosophila* genital tract regulate female reproductive behavior. Neuron 61: 511–518.

Hayes, R. O., 1953 Determination of a Physiological Saline Solution for *Aedes aegypti* (L.). Journal of Economic Entomology 46: 624–627.

Helinski, M. E., P. Deewatthanawong, L. K. Sirot, M. F. Wolfner and L. C. Harrington, 2012 Duration and dose-dependency of female sexual receptivity responses to seminal fluid proteins in *Aedes albopictus* and *Ae. aegypti* mosquitoes. J Insect Physiol 58: 1307–1313.

Helinski, M. E., and L. C. Harrington, 2011 Male mating history and body size influence female fecundity and longevity of the dengue vector *Aedes aegypti*. J Med Entomol 48: 202–211.

Hiss, E. A., and M. S. Fuchs, 1972 The effect of matrone on oviposition in the mosquito, *Aedes Aegypti*. J Insect Physiol 18: 2217–2227.

Hua, Y. J., Y. Tanaka, K. Nakamura, M. Sakakibara, S. Nagata et al., 1999 Identification of a prothoracicostatic peptide in the larval brain of the silkworm, *Bombyx mori*. J Biol Chem 274: 31169–31173.

Hull, J. J., and C. S. Brent, 2014 Identification and characterization of a sex peptide receptor-like transcript from the western tarnished plant bug *Lygus hesperus*. Insect Mol Biol 23: 301–319.

Hussain, A., H. K. Ucpunar, M. Zhang, L. F. Loschek and I. C. Grunwald Kadow, 2016 Neuropeptides Modulate Female Chemosensory Processing upon Mating in *Drosophila*. PLoS Biol 14: e1002455.

Judson, C. L., 1967 Feeding and oviposition behavior in the mosquito *Aedes aegypti* (L.). I. Preliminary studies of physiological control mechanisms. Biol Bull 133: 369–378.

Kim, Y. J., K. Bartalska, N. Audsley, N. Yamanaka, N. Yapici et al., 2010 MIPs are ancestral ligands for the sex peptide receptor. Proc Natl Acad Sci U S A 107: 6520–6525.

Kim, Y. J., D. Zitnan, K. H. Cho, D. A. Schooley, A. Mizoguchi et al., 2006a Central peptidergic ensembles associated with organization of an innate behavior. Proc Natl Acad Sci U S A 103: 14211–14216.

Kim, Y. J., D. Zitnan, C. G. Galizia, K. H. Cho and M. E. Adams, 2006b A command chemical triggers an innate behavior by sequential activation of multiple peptidergic ensembles. Curr Biol 16: 1395–1407.

Kistler, K. E., L. B. Vosshall and B. J. Matthews, 2015 Genome engineering with CRISPR-Cas9 in the mosquito *Aedes aegypti*. Cell Rep 11: 51–60.

Labun, K., T. G. Montague, J. A. Gagnon, S. B. Thyme and E. Valen, 2016 CHOPCHOP v2: a web tool for the next generation of CRISPR genome engineering. Nucleic Acids Res 44: W272–276.

Lang, C. A., 1956 The influence of mating on egg production by *Aedes aegypti*. Am J Trop Med Hyg 5: 909–914.

Leahy, M. G., 1967 Non-specificity of the male factor enhancing egg-laying in Diptera. Journal of Insect Physiology 13: 1283–1292.

Leahy, M. G., and G. B. Craig, Jr., 1965 Accessory gland substance as a stimulant for oviposition in *Aedes aegypti* and *A. albopictus*. Mosquito News 25: 448–452.

Lee, J. H., N. R. Lee, D. H. Kim and Y. J. Kim, 2020 Molecular characterization of ligand selectivity of the sex peptide receptors of *Drosophila melanogaster* and *Aedes aegypti*. Insect Biochem Mol Biol 127: 103472.

Li, C., J.-F. Yu, Q. Lu, J. Xu, J.-H. Liu et al., 2014 Molecular characterization and functional analysis of a putative sex-peptide receptor in the tobacco cutworm Spodoptera litura(Fabricius, 1775) (Lepidoptera: Noctuidae). Austral Entomology 53: 424–431.

Li, M., M. Bui, T. Yang, C. S. Bowman, B. J. White et al., 2017 Germline Cas9 expression yields highly efficient genome engineering in a major worldwide disease vector, *Aedes aegypti*. Proc Natl Acad Sci U S A 114: E10540–E10549.

Liu, H., and E. Kubli, 2003 Sex-peptide is the molecular basis of the sperm effect in *Drosophila melanogaster*. Proc Natl Acad Sci U S A 100: 9929–9933.

Liu, S., B. Li, W. Liu, Y. Liu, B. Ren et al., 2021 Sex peptide receptor mediates the post-mating switch in *Helicoverpa armigera* (Lepidoptera: Noctuidae) female reproductive behavior. Pest Manag Sci 77: 3427–3435.

Lorenz, M. W., R. Kellner and K. H. Hoffmann, 1995 A family of neuropeptides that inhibit juvenile hormone biosynthesis in the cricket, *Gryllus bimaculatus*. J Biol Chem 270: 21103–21108.

Lubawy, J., P. Marciniak and G. Rosinski, 2020 Identification, Localization in the Central Nervous System and Novel Myostimulatory Effect of Allatostatins in *Tenebrio* molitor Beetle. Int J Mol Sci 21.

Matsumoto, K., Y. Suetsugu, Y. Tanaka, T. Kotaki, S. G. Goto et al., 2017 Identification of allatostatins in the brown-winged green bug *Plautia stali*. J Insect Physiol 96: 21–28.

Matthews, B. J., C. S. McBride, M. DeGennaro, O. Despo and L. B. Vosshall, 2016 The neurotranscriptome of the *Aedes aegypti* mosquito. BMC Genomics 17: 32.

McGeary, M. K., and G. D. Findlay, 2020 Molecular evolution of the sex peptide network in *Drosophila*. J Evol Biol 33: 629–641.

McMeniman, C. J., R. A. Corfas, B. J. Matthews, S. A. Ritchie and L. B. Vosshall, 2014 Multimodal integration of carbon dioxide and other sensory cues drives mosquito attraction to humans. Cell 156: 1060–1071.

Min, S., H. S. Chae, Y. H. Jang, S. Choi, S. Lee et al., 2016 Identification of a Peptidergic Pathway Critical to Satiety Responses in *Drosophila*. Curr Biol 26: 814–820.

Naccarati, C., N. Audsley, J. N. Keen, J. H. Kim, G. J. Howell et al., 2012 The host-seeking inhibitory peptide, Aea-HP-1, is made in the male accessory gland and transferred to the female during copulation. Peptides 34: 150–157.

Nasci, R. S., 1990 Relationship of wing length to adult dry weight in several mosquito species (Diptera: Culicidae). J Med Entomol 27: 716–719.

Nene, V., J. R. Wortman, D. Lawson, B. Haas, C. Kodira et al., 2007 Genome sequence of *Aedes aegypti*, a major arbovirus vector. Science 316: 1718–1723.

Offermanns, S., and M. I. Simon, 1995 Gα15 and Gα16 Couple a Wide Variety of Receptors to Phospholipase C (∗). Journal of Biological Chemistry 270: 15175–15180.

Oh, Y., S. E. Yoon, Q. Zhang, H. S. Chae, I. Daubnerova et al., 2014 A homeostatic sleep-stabilizing pathway in *Drosophila* composed of the sex peptide receptor and its ligand, the myoinhibitory peptide. PLoS Biol 12: e1001974.

Okamoto, N., and A. Watanabe, 2022 Interorgan communication through peripherally derived peptide hormones in *Drosophila*. Fly (Austin) 16: 152–176.

Paluzzi, J. P., A. S. Haddad, L. Sedra, I. Orchard and A. B. Lange, 2015 Functional characterization and expression analysis of the myoinhibiting peptide receptor in the Chagas disease vector, *Rhodnius prolixus*. Mol Cell Endocrinol 399: 143–153.

Poels, J., T. Van Loy, H. P. Vandersmissen, B. Van Hiel, S. Van Soest et al., 2010 Myoinhibiting peptides are the ancestral ligands of the promiscuous *Drosophila* sex peptide receptor. Cell Mol Life Sci 67: 3511–3522.

Predel, R., S. Neupert, S. F. Garczynski, J. W. Crim, M. R. Brown et al., 2010 Neuropeptidomics of the mosquito *Aedes aegypti*. J Proteome Res 9: 2006–2015.

Santos, J. G., M. Vomel, R. Struck, U. Homberg, D. R. Nassel et al., 2007 Neuroarchitecture of peptidergic systems in the larval ventral ganglion of *Drosophila melanogaster*. PLoS One 2: e695.

Scheunemann, L., A. Lampin-Saint-Amaux, J. Schor and T. Preat, 2019 A sperm peptide enhances long-term memory in female *Drosophila*. Sci Adv 5: eaax3432.

Schmidt, T., Y. Choffat, S. Klauser and E. Kubli, 1993 The *Drosophila melanogaster* sex-peptide: A molecular analysis of structure-function relationships. Journal of Insect Physiology 39: 361–368.

Schoofs, L., G. M. Holman, T. K. Hayes, R. J. Nachman and A. De Loof, 1991 Isolation, primary structure, and synthesis of locustapyrokinin: a myotropic peptide of *Locusta migratoria*. Gen Comp Endocrinol 81: 97–104.

Siju, K. P., A. Reifenrath, H. Scheiblich, S. Neupert, R. Predel et al., 2014 Neuropeptides in the antennal lobe of the yellow fever mosquito, *Aedes aegypti*. J Comp Neurol 522: 592–608.

Smith, L. B., S. Kasai and J. G. Scott, 2018 Voltage-sensitive sodium channel mutations S989P + V1016G in *Aedes aegypti* confer variable resistance to pyrethroids, DDT and oxadiazines. Pest Manag Sci 74: 737–745.

Smith, R. C., M. F. Walter, R. H. Hice, D. A. O’Brochta and P. W. Atkinson, 2007 Testis-specific expression of the beta2 tubulin promoter of *Aedes aegypti* and its application as a genetic sex-separation marker. Insect Mol Biol 16: 61–71.

Swanson, W. J., and V. D. Vacquier, 2002 The rapid evolution of reproductive proteins. Nat Rev Genet 3: 137–144.

Tsuda, M., and T. Aigaki, 2016 Evolution of sex-peptide in *Drosophila*. Fly (Austin) 10: 172–177.

Tsuda, M., J. B. Peyre, T. Asano and T. Aigaki, 2015 Visualizing Molecular Functions and Cross-Species Activity of Sex-Peptide in *Drosophila*. Genetics 200: 1161–1169.

Villarreal, S. M., S. Pitcher, M. E. H. Helinski, L. Johnson, M. F. Wolfner et al., 2018 Male contributions during mating increase female survival in the disease vector mosquito *Aedes aegypti*. J Insect Physiol 108: 1–9.

Wang, F., K. Wang, N. Forknall, C. Patrick, T. Yang et al., 2020 Neural circuitry linking mating and egg laying in *Drosophila* females. Nature 579: 101–105.

Wang, K., F. Wang, N. Forknall, T. Yang, C. Patrick et al., 2021 Neural circuit mechanisms of sexual receptivity in *Drosophila* females. Nature 589: 577–581.

White, M. A., A. Bonfini, M. F. Wolfner and N. Buchon, 2021 *Drosophila melanogaster* sex peptide regulates mated female midgut morphology and physiology. Proc Natl Acad Sci U S A 118.

Wigby, S., N. C. Brown, S. E. Allen, S. Misra, J. L. Sitnik et al., 2020 The *Drosophila* seminal proteome and its role in postcopulatory sexual selection. Philos Trans R Soc Lond B Biol Sci 375: 20200072.

Wohl, M. P., and C. J. McMeniman, 2023a Batch Rearing *Aedes aegypti*. Cold Spring Harb Protoc 2023: pdb prot108017.

Wohl, M. P., and C. J. McMeniman, 2023b Single-Pair and Small-Group Crosses of *Aedes aegypti*. Cold Spring Harb Protoc 2023: pdb prot108018.

Yang, C. H., S. Rumpf, Y. Xiang, M. D. Gordon, W. Song et al., 2009 Control of the postmating behavioral switch in *Drosophila* females by internal sensory neurons. Neuron 61: 519–526.

Yapici, N., Y. J. Kim, C. Ribeiro and B. J. Dickson, 2008 A receptor that mediates the post-mating switch in *Drosophila* reproductive behaviour. Nature 451: 33–37.

Zheng, W., Y. Liu, W. Zheng, Y. Xiao and H. Zhang, 2015 Influence of the silencing sex-peptide receptor on *Bactrocera dorsalis* adults and offspring by feeding with ds-spr. Journal of Asia-Pacific Entomology 18: 477–481.

